# Non-Invasive Mechanical-Functional Analysis of Individual Liver Mitochondria by Atomic Force Microscopy

**DOI:** 10.1101/2025.06.01.657242

**Authors:** Ekaterina O. Zorikova, Sabita Chourasia, Irit Rosenhek-Goldian, Sidney R. Cohen, Semen V. Nesterov, Atan Gross

**Author notes:** Corresponding author: Atan Gross, PhD, Department of Immunology and Regenerative Biology Weizmann Institute of Science, Rehovot Israel 76100.

## Abstract

Mitochondria play a pivotal role in energy production, signaling, and apoptosis. Yet, probing their functional state at the single-organelle level without invasive labels remains a major challenge. Here, we introduce a novel, label-free approach that leverages Atomic Force Microscopy (AFM) beyond its traditional imaging role, transforming it into a powerful tool for functional analysis of individual, isolated mitochondria. By immobilizing mouse liver mitochondria on polylysine-coated mica, we achieved nanoscale resolution of mitochondrial mechanical properties including height, height fluctuation power spectra, and Young’s modulus, under different respiratory states. Strikingly, fluctuations in mitochondrial height fluctuations below 20 Hz showed robust correlation with the mitochondria membrane potential (ΔΨ_m_), a cornerstone of mitochondrial function. This relationship allows AFM to sensitively detect changes in the mitochondria bioenergetic status. Applying this method to mitochondria from liver-specific MTCH2 liver-conditional knockout mice, a model of mitochondrial malfunction, we confirmed AFM’s diagnostic potential. The technique reliably distinguished malfunctional mitochondria, mirroring and adding new insights beyond conventional fluorescence assays. By bridging nanomechanics and mitochondrial bioenergetics, this approach paves the way for non-invasive, high-resolution diagnostics at the single-organelle level, holding promise to monitor the actual functional state of mitochondria in clinical settings.

## Introduction

Mitochondria are organelles of eukaryotic cells that play a crucial role in energy production and regulating cellular metabolism. They function as the biochemical powerhouses of the cell, generating adenosine triphosphate (ATP) through the process of oxidative phosphorylation. The electron transport chain (ETC), which consists of five main complexes, is localized in the mitochondrial inner membrane and generates an electrochemical potential that drives ATP-synthase ^1^. Assessing the function of individual ETC complexes is crucial in biomedical research, as mitochondria play a pivotal role in cellular energy metabolism, pro-inflammatory signaling, and apoptosis ^2–4^. Mitochondrial dysfunction is implicated in a wide range of diseases and pathological conditions. Various methods exist for routine mitochondrial function analysis, each with its own advantages and limitations.

Fluorescent techniques are widely used to assess mitochondrial properties in intact cells as well as in isolated mitochondria. These techniques provide measurements of ΔΨ_m_, reactive oxygen species (ROS) production, and mitochondrial network integrity^5–7^. Most accurate diagnostics usually are carried out on isolated organelles. These include, but are not limited to, different ΔΨ_m_- and ROS-sensitive fluorescent dyes, electron microscopy, selective electrodes for oxygen, pH, TPP^+^ (ΔΨ_m_ measurement), and Ca^2+^ ^8–11^. However, in contrast to structural studies, there is a lack of generally accepted techniques which can monitor the functioning of single mitochondria. The activity of specific ion channels and exchangers can be studied using the patch-clamp technique ^12–15^. This method is quite challenging due to the small size of mitochondria and is limited to measuring electrical properties.

A popular method for studying parameters of single cells and more rarely of isolated mitochondria is flow cytometry, including Fluorescence-activated Cell Sorting (FACS)^16^. The advantage of FACS is the ability to analyze many objects with simultaneous assessment of several characteristics, such as ΔΨ_m_, oxidative stress and size ^17,18^. However, the method also has its drawbacks because the potential-sensitive dyes are accumulated in the inner membrane or matrix and can affect mitochondrial function. For example, rhodamine derivatives can induce photo-toxicity ^19^, and safranin may inhibit complex I in neuronal cells^20^. Low concentrations of dyes are used whenever possible, but the sensitivity to side effects of mitochondrial dyes from different tissues and organisms, in normal and pathological conditions, varies greatly and unpredictably, which can result in misleading results. The artifacts can also be induced by the movement of dyes between quenched (hydrophobic) and unquenched (hydrophilic) compartments ^21^. It is also important to note that the FACS method does not have high-resolution capabilities. Therefore, mitochondria are visualized as practically point objects close in size to the fluorescence wavelength, leading to large standard deviation and limiting the analysis to large populations, making this method not much better than ensemble measurement of a suspension.

AFM is a powerful method for studying biological objects ^22^. Typically, a tens of micrometer-sized cantilever with a nanometer-sized, sharp tip is used for biological applications ^23^. The tip can scan the surface, line by line, and track its morphology. AFM visualization can be performed in very gentle modes to avoid damage to soft samples, in both air and liquid environments ^24–26^. AFM can be used to measure not only sizes of nanoscale objects and surface roughness, but also elastic properties of the sample ^27–29^. Here we apply AFM for imaging, mechanical analysis, and frequency-dependent height fluctuations to obtain multi-functional single organelle analysis in a liquid environment without chemical modifications of the mitochondria.

Specifically, our proof-of-principle experiments demonstrate that AFM measurements can distinguish different mitochondrial functional states which are characterized by variations in membrane potential. Moreover, AFM enables the detection of mitochondrial abnormalities, as demonstrated in our tests using mitochondria prepared from MTCH2 liver-conditional knockout mice. The universal mechanical-functional approach we have developed could serve as a foundation for novel diagnostic methods, enabling the detection of mitochondrial malfunctions in clinical settings.

## Results

### Mitochondria height fluctuations measured by AFM show excellent correlation with ΔΨ_m_ measured by potentiometric fluorescent dyes

In this study, we used isolated mouse liver mitochondria (MLM), a well-established model system for bioenergetic research^46^. A representative AFM image of MLM obtained in liquid tapping mode is shown in Figure 1A. The purity and integrity of the MLM preparations were confirmed via confocal microscopy using the ΔΨ_m_-sensitive dye tetramethylrhodamine methyl ester (TMRM), alongside carbonyl cyanide m-chlorophenyl hydrazone (CCCP), a protonophore that dissipates ΔΨ_m_ and serves as a control for mitochondrial dysfunction (Figures S1A, B in Supporting Information).

**Figure 1.**
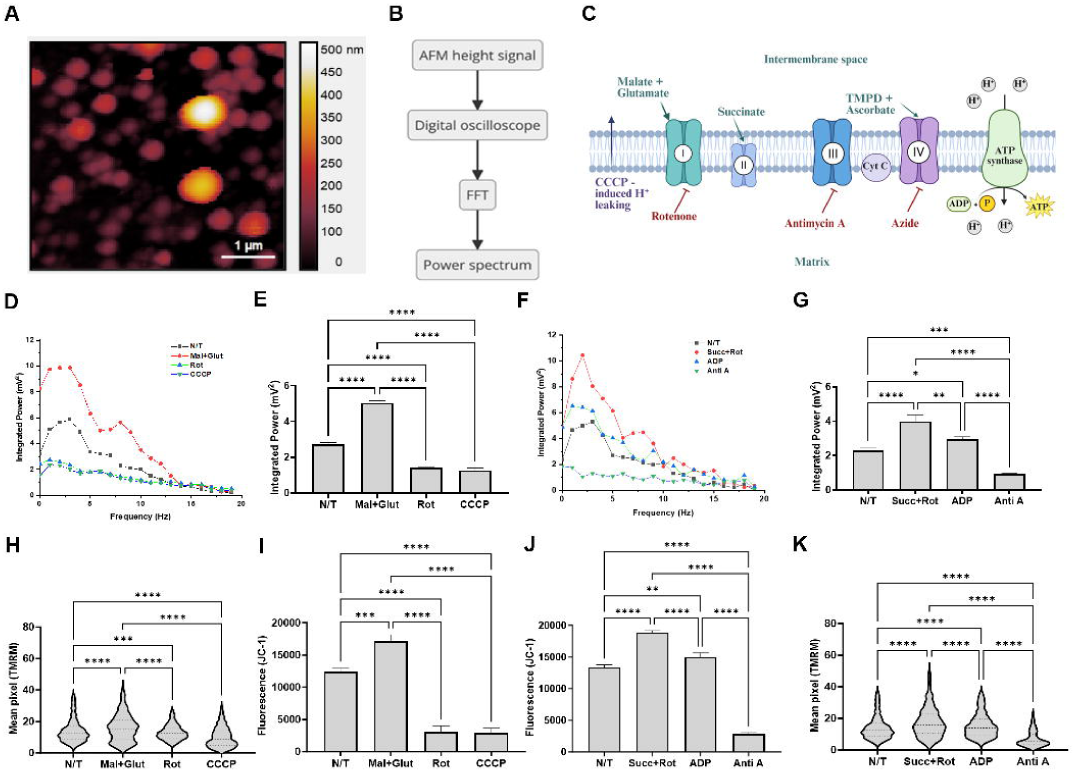
Correlation of mitochondrial membrane potential (ΔΨm) with the power spectrum of mitochondria’s height fluctuations (A) Representative AFM topography image of isolated mouse liver mitochondria (MLM) adsorbed onto poly-L-lysine–coated mica, acquired in liquid tapping mode. (B) Workflow for extracting power spectra from AFM height signals, including digital acquisition, fast Fourier transform (FFT), and spectral analysis. (C) Diagram of the electron transport chain (ETC) illustrating the key respiratory substrates (green) and inhibitors (red) used in this study. (D) Smoothed power spectra of mitochondrial height fluctuations under Complex I-linked conditions (malate + glutamate, rotenone, CCCP). Power spectra are normalized to allow visualization of power enhancements at different frequency ranges (see methods). (E) Quantification of integrated power (0–20 Hz range) for the conditions shown in panel D. (F) Smoothed power spectra under Complex II-linked conditions (succinate + rotenone, ADP, antimycin A). (G) Quantification of integrated power (0–20 Hz range) for the conditions shown in panel F. (H) Quantification of ΔΨm using TMRM fluorescence measured by flow cytometry. (I) Quantification of ΔΨm using JC-1 fluorescence measured by a plate reader for conditions in panel D. (J) Quantification of ΔΨm using JC-1 fluorescence for conditions in panel F. (K) Quantification of ΔΨm using TMRM fluorescence for conditions in panel F. All data represent mean ± SEM; n = 30 individual mitochondria per condition (AFM). Statistical analysis was performed using one-way ANOVA with multiple comparisons: ns = not significant; *p ≤ 0.05, **p ≤ 0.01, ***p ≤ 0.001, ****p ≤ 0.0001.

Additional confirmation was obtained through transmission electron microscopy (Figure S1C in Supporting Information.) and AFM images of dried mitochondria, revealing characteristic cristae structures (Figures S1D, E in Supporting Information). We also measured the oxygen consumption rate (OCR) using seahorse analysis, and as anticipated, the addition of CCCP or ADP (an activator of phosphorylation) increased OCR, whereas rotenone and antimycin A, inhibitors of complex I and III respectively, markedly suppressed OCR (Figures S1F, G in Supporting Information).

Using topographic images such as that shown in Figure 1A as a guide, the AFM probe was positioned on the surface of individual mitochondria, and height fluctuations were monitored over time under feedback control. The analysis algorithm is illustrated in Figure 1B, where the AFM height signal was processed through a digital oscilloscope and subjected to fast Fourier transform (FFT) to yield a power spectrum. The low-frequency range (0–20 Hz) of the power spectrum was analyzed to assess mitochondrial responses to different functional states.

The power spectra exhibited distinct and characteristic responses to various respiratory substrates and inhibitors, consistent with their known activities (Figure 1C). Specifically, responses to malate + glutamate (complex I substrates), rotenone (a complex I inhibitor), and CCCP (an uncoupler) are shown in Figures 1D, E. Further responses to succinate (a complex II substrate with rotenone to inhibit complex I), ADP (an activator of phosphorylation), and antimycin A (a complex III inhibitor) are presented in Figures 1F, G. Additional analyses of TMPD + ascorbate (complex IV substrates) and azide (a complex V inhibitor) are shown in Figures S1H, I in Supporting Information, and spectra for the additives alone with no mitochondria are presented in Figure S1J in Supporting Information.

To validate the AFM-based assessment of mitochondrial function, we compared the integrated power from AFM spectra with measurements of ΔΨ_m_ obtained using two potentiometric fluorescent dyes: JC-1 analyzed using a fluorescence plate reader, and TMRM analyzed by flow cytometry. Remarkably, ΔΨ_m_ values obtained via both JC-1 and TMRM exhibited excellent correlation with the integrated power across the tested conditions described above (compare Figure 1E with 1H, I, Figure 1G with 1J, K, and Figure S1I with S1K, L).

Across all experimental modalities, the addition of respiratory substrates led to an increase in integrated power (AFM) and in ΔΨ_m_ (TMRM and JC-1), whereas inhibitors of the respective complexes induced a reduction in these parameters. Notably, the addition of ADP, which stimulates phosphorylation, resulted in a moderate decline in ΔΨ_m_ (by approximately 10–20%), consistent across all three methods. Conversely, treatment with CCCP caused an almost complete collapse of ΔΨ_m_.

While all three methods demonstrated qualitative agreement, the quantitative magnitudes of the changes varied, indicating that calibration would be required for direct quantitative comparisons. Nonetheless, for diagnostic or functional assessment purposes, qualitative or semi-quantitative analyses (e.g., expressed as a percentage of control) are often sufficient. These results demonstrate that AFM-based assessment of integrated power provides a reliable and comparable measure of ΔΨ_m_, on par with established fluorescence-based techniques. The details of the frequency response are depicted qualitatively in the power spectra such as Figures 1D and E, and the total integrated power up to 20 Hz shown quantitatively in bar graphs, such as Figures 1 E and G.

### Mitochondrial elastic modulus and height fluctuations are inversely correlated

A distinctive advantage of AFM over most other methods is its ability to simultaneously analyze surface morphology and elastic properties. By imaging a field of view containing multiple mitochondria, as shown in Figure 1A, AFM allows for precise estimation of mitochondrial size and height. Elastic properties are measured simultaneously to the topography, allowing direct correlation between them.

Figure 2 presents a comprehensive analysis of the height and elastic properties of mitochondria. Figures 2A and B show maps of the Young’s modulus. In addition to providing information on mechanical properties of the mitochondria, this characteristic can also be useful for distinguishing mitochondria from various polymer aggregates (protein aggregates, lipoprotein complexes, vesicles, etc.), which have similar sizes but different rigidities.

**Figure 2.**
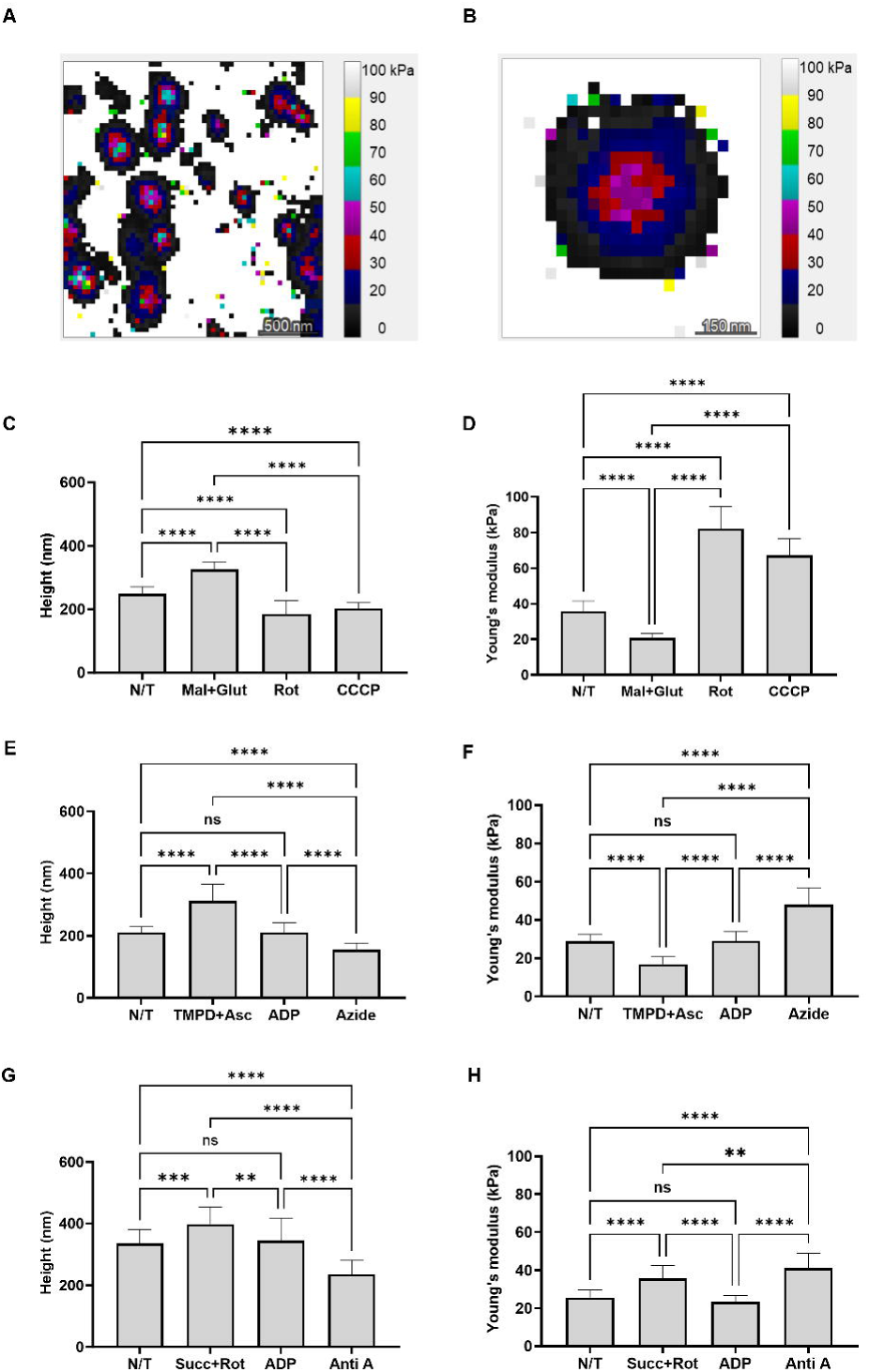
Mitochondrial height and stiffness (Young’s modulus) under different respiratory conditions assessed by AFM (A) Representative AFM Young’s modulus map of multiple mitochondria adhered to a poly-L-lysine–coated mica surface. (B) High-resolution Young’s modulus map of a single mitochondrion showing spatial stiffness distribution. (C) Height measurements of individual mitochondria under Complex I-linked conditions (non-treated, malate + glutamate, rotenone, CCCP). (D) Young’s modulus of mitochondria under the same conditions as in panel C. (E) Height measurements under Complex IV-linked conditions (TMPD + ascorbate, ADP, azide). (F) Young’s modulus under the same conditions as in panel E. (G) Height measurements under Complex II-linked conditions (succinate + rotenone, ADP, antimycin A). (H) Young’s modulus under the same conditions as in panel G. All data represent mean ± SEM; n = 30 individual mitochondria per condition. Statistical significance was evaluated by one-way ANOVA with multiple comparisons: ns = not significant; *p ≤ 0.05, **p ≤ 0.01, ***p ≤ 0.001, ****p ≤ 0.0001.

The changes in height can be directly compared with simultaneously obtained elastic properties of mitochondria in different functional states as described in Figure 1. Activation of complex I with glutamate and malate resulted in an increase in mitochondrial height (Figure 2C) whereas the elastic modulus/stiffness was decreased (Figure 2D). A similar inverse correlation was obtained following activation of complex IV with TMPD and ascorbate (Figures 2E, F). On the other hand, a decrease in ΔΨ_m_ caused by the protonophore CCCP or each of the respiratory chain inhibitors, rotenone, antimycin A, azide, led to a reduction in mitochondrial height (Figures 2C, E, respectively) whereas the elastic modulus/stiffness was increased (Figure 2D, F, respectively). As one might expect from the results described above, the addition of succinate with rotenone (complex II substrate with complex I inhibitor) resulted in a “hybrid” behavior – an increase in both mitochondria height and elastic modulus/stiffness (Figures 2 G, H, respectively).

### Integrated power does not correlate with mitochondrial swelling

The straight correlation of integrated power with mitochondrial height raises the question of whether mitochondrial height depends on ΔΨ_m_ or that only fluctuation amplitude has such dependence. It is known from transmission electron microscopy experiments that an increase in ΔΨ_m_ can cause swelling of the mitochondrial matrix^30^. The results showing changes in the size and elastic properties in different functional states of mitochondria confirm that there are indeed changes in the mitochondrial physical properties following alterations in its functional state. Since an increase in potential across the inner mitochondrial membrane (IMM) leads to an influx of osmotically active cations into the matrix (mainly K+ ions), it was necessary to consider whether AFM registers changes related to this process. To unambiguously identify conditions for which it is correct to interpret changes in integrated power as changes in ΔΨm, it was necessary to check how this and other parameters measured on AFM are related to changes in the mitochondrial matrix volume.

The integrated power of mitochondrial membrane fluctuations decreased significantly upon valinomycin treatment (Figure 3A), indicating a drop in ΔΨ_m_. This was accompanied by an increase in mitochondrial height (Figure 3B), suggesting matrix swelling caused by osmotic imbalance due to enhanced KL influx. Swelling also led to mechanical softening of mitochondria, as shown by a decrease in Young’s modulus (Figure 3C). These structural and mechanical changes were accompanied by a loss of membrane potential, confirmed by reduced TMRM fluorescence (Figure 3D) and JC-1 aggregate fluorescence (Figure 3E). Subsequent treatment with CCCP caused a further decline in ΔΨ_m_ across all measurements.

**Figure 3.**
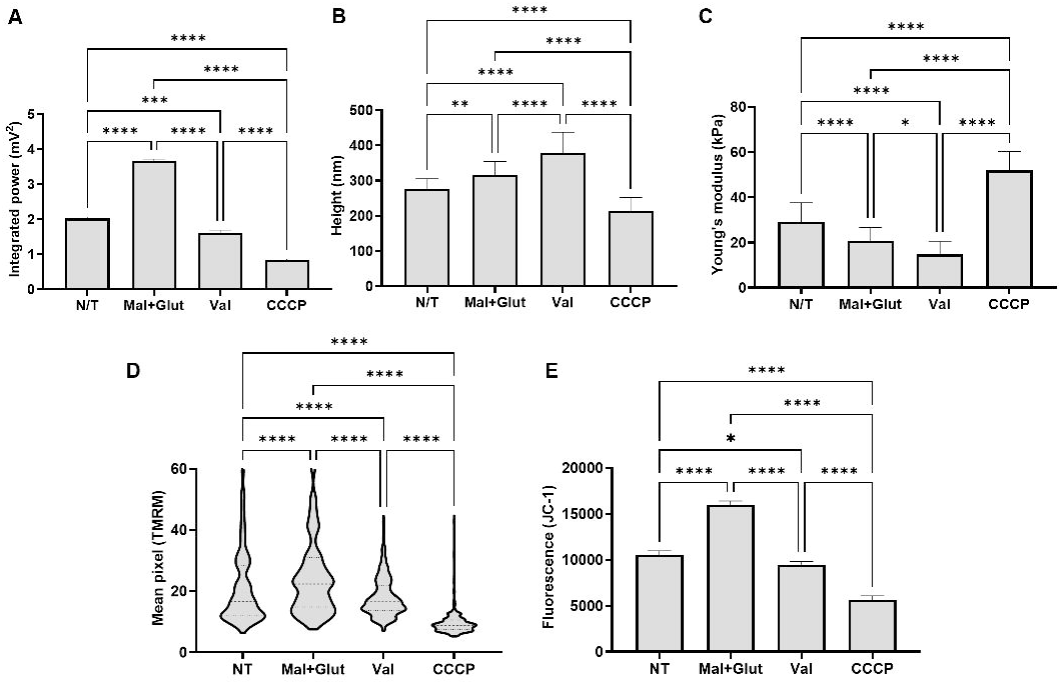
Effects of mitochondrial matrix swelling on AFM and fluorescence-based measurements (A) Integrated power of mitochondrial height fluctuations under different conditions (non-treated, malate + glutamate, valinomycin, CCCP), showing a marked decrease in power with valinomycin, indicative of reduced membrane potential. (B) Height distribution of individual mitochondria demonstrating matrix swelling in the presence of valinomycin, likely due to osmotic imbalance following potassium influx. (C) Young’s modulus distribution showing mechanical softening of mitochondria under valinomycin and CCCP, reflecting changes in stiffness. (D) Mitochondrial membrane potential measured via TMRM fluorescence using flow cytometry. (E) JC-1 fluorescence-based assessment of ΔΨm using a plate reader. All data represent mean ± SEM; n = 30 mitochondria per condition for AFM. Statistical significance was assessed by one-way ANOVA with multiple comparisons: ns = not significant; *p ≤ 0.05, **p ≤ 0.01, ***p ≤ 0.001, ****p ≤ 0.0001.

Pearson correlation coefficients reveal that neither mitochondrial height nor Young’s modulus strongly correlate with changes in integrated power across the conditions tested (all |r| < 0.3). This suggests that the changes in mechanical properties and swelling do not follow the energetic shifts due to changes in ΔΨ_m_. The inverse response in comparison to Figures 1 and 2 between integrated power and height changes in these experiments shows that integrated power is independent of height change. Thus, the correlation of integrated power with ΔΨ_m_, but not with swelling, is confirmed.

### AFM analysis can detect functional defects in mitochondria

To assess AFM’s ability to detect functional defects in mitochondria, we examined MLM isolated from MTCH2 liver-conditional knockout mice. MTCH2 is a key regulator of apoptosis, metabolism, and mitochondrial dynamics ^31–33^. Its precise mechanism of action remains largely unknown, but AFM analysis may provide further insights into MTCH2’s function by revealing mechanical alterations in mitochondria lacking this protein.

AFM analysis indeed revealed changes in the mechanical and bioenergetic properties of MTCH2 knockout (MKO) mitochondria compared to wild type (WT) mitochondria. Specifically, MKO mitochondria exhibited increased integrated power (Figure 4A), which is again correlated with an elevation in ΔΨ_m_ (Figures 4B, C). In addition, MKO mitochondria displayed increased modulus/stiffness (Figure 4D), along with a decrease in mitochondrial height (Figure 4E).

**Figure 4.**
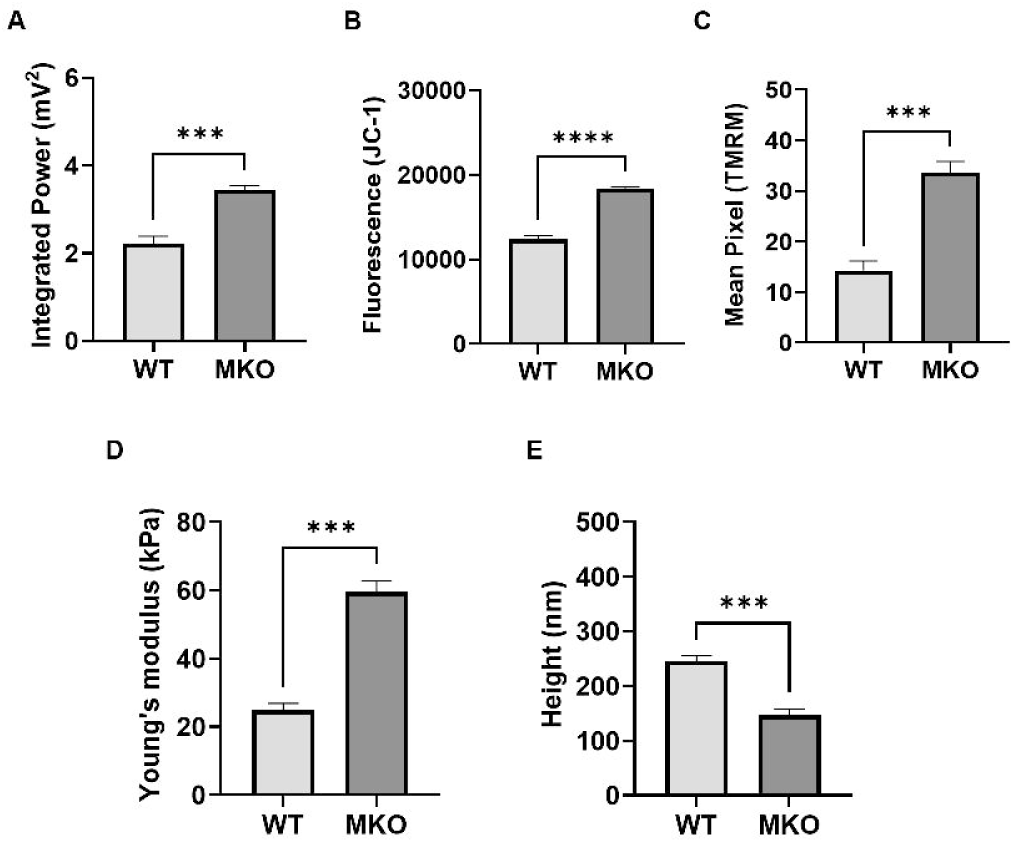
Biophysical and functional differences between mitochondria from wild-type (WT) and MTCH2 liver-conditional knockout (MKO) mice under non-treated conditions (A) Integrated power of height fluctuations measured by AFM, reflecting mitochondrial membrane potential. (B) JC-1 fluorescence intensity measured by a plate reader as an indicator of ΔΨm. (C) TMRM mean pixel intensity measured by flow cytometry to assess ΔΨm. (D) Young’s modulus determined by AFM force spectroscopy, indicating mitochondrial stiffness. (E) Mitochondrial height measured by AFM topography. MKO mitochondria show significantly higher membrane potential (A–C), increased stiffness (D), and reduced height (E) compared to WT controls. Data represent mean ± SEM from three biological replicates (10 mitochondria per replicate). Statistical analysis was performed using one-way ANOVA with multiple comparisons: ns = not significant; *p ≤ 0.05, **p ≤ 0.01, ***p ≤ 0.001, ****p ≤ 0.0001.

The increased modulus/stiffness observed in the MKO MLM mirrors the increase observed with WT MLM treated with either mitochondria uncouplers or respiratory complex inhibitors (Figure 2), suggesting that MTCH2 deletion results in mitochondria uncoupling and/or inhibition of one of the respiratory complexes. This is a novel finding we did not detect using standard techniques. On the other hand, MKO MLM show an increase in integrated power, which is the opposite of the decrease seen after treatment with either mitochondria uncouplers or respiratory complex inhibitors (Figure 1), suggesting that MTCH2 deletion increases mitochondrial activity, as we have previously reported in several papers ^31–33^.

## Discussion

By tracking the height fluctuations (“noise”) of the organelle with time, AFM is able to monitor the dynamics of energy flow. AFM was first used to monitor activity in a biological system by looking at height fluctuations on a protein confined to a surface upon exposure to its enzymatic substrate ^39^. Mitochondria represent a more complex system of study, requiring additional input. Accordingly, the AFM method employed here measured several characteristics such as morphology and elastic modulus of the structures, as well as fully exploiting the rich information available from the noise spectra.

To understand how this information can be related to mitochondrial function, we need to dissect the components of the noise signal. Power spectra, such as those shown in Figure 1D, F shows an overall trend of decreasing power with frequency, with enhancement at two different frequency ranges (2-4 Hz and 6-9 Hz) superimposed on this background. Both the broad envelope and the prominent peaks are of interest. The background profile can be described by frequency dependence of 1/f^α^ for mitochondria both untreated, and under activation or inhibition. Interestingly, the control traces and more importantly the inhibited mitochondria, show a flat or nearly flat trend (α≈0). Living organisms are inherently noisy, and this noise is important in maintaining homeostasis ^40^.

Noise plays an important part in harnessing various physiological regulatory processes occurring at different frequencies. Frequency behavior where the 1/f^α^ noise has an exponent α close to 1 is termed “pink” noise and is characteristic of viable systems. This behavior is found, for instance, in the human heartbeat. Previous studies have shown that most viable systems have an exponent close to 1. An exponent of zero, otherwise known as “white noise” would result in a “flat” power spectrum with equal energy at all frequencies and is indicative of an aging or unhealthy organism^40^. This α≈0 behavior, indicated by weak dependence in the log-log plot of power vs. frequency (poor correlation between log power and log frequency) is seen after inhibitor additions such as rotenone. This is also the case for control on the mica substrate. These are shown in Figures S2A, B, respectively. Interestingly, untreated mitochondria also give intermediate values, around α=0.6, but enhancement with Malate/Glutamate gives an exponent > 1 (see Figures S2C, D).

The second component of these spectra is the distinct enhancement seen at certain frequency ranges. The enhancement over the range 2-4 Hz is seen both for untreated mitochondria and for all substrates that enhance the activity. It is never seen under inhibition. This frequency was observed previously on ensemble systems. One of these works employed dynamic phase microscopy and associated it with action of the ATPase pump ^41^. A second work, in which monitored noise in fluctuations of an AFM cantilever upon which mitochondria were absorbed, found a distinct peak at 1-3 Hz. As in our work, these oscillations were suppressed when inhibited by rotenone ^42^. Since these independent data were taken from mitochondria extracted from two diverse sources of human cells, that frequency range seems to be a general characteristic of mitochondria. The distinguishing aspect of this work is that we measure single organelles and can correlate the size, modulus, and noise function with the mitochondrial state. The second peak we observed at 6-8 Hz only appears under activation and seems to be a new observation here as it is not mentioned in previous literature.

Integrating the power spectrum over the entire frequency range gives integrated power i.e., RMS noise. These data are plotted in Figure 1, showing remarkable correlation of this quantity with the ΔΨ_m_ as recorded by fluorescent techniques (FACS and plate). Whereas this quantity does not carry information on the finer features discussed above, it forms an important basis for the strength of this method in monitoring the membrane activity nondestructively.

The ability of AFM to detect changes in mitochondrial properties under pathological conditions, as demonstrated with MTCH2 KO (MKO) MLM, underscores its diagnostic potential. MKO MLM exhibited increased integrated power/ΔΨ_m_, increased modulus/stiffness, and decreased height. These findings suggest that AFM could be employed to identify mitochondrial malfunctions associated with metabolic disorders or degenerative diseases. In particular, the observed mechanical remodeling in MKO MLM could serve as a biomarker for early detection of mitochondrial pathologies, even when traditional methods fail to reveal significant structural changes.

## Conclusion

Our data underscores the necessity of combining various techniques for a comprehensive understanding of the biophysical and biochemical processes occurring in mitochondria. The data also prove our ability to monitor single mitochondria function with the AFM probe. We hope that the developed methodology will provide more fundamentally relevant data on mitochondria, as well as improve the quality of diagnosis of mitochondria-associated diseases and make these studies more widely accessible.

## Materials and methods

### Animals and Ethical Compliance

All animal work was conducted in accordance with European guidelines for the care and use of laboratory animals (Directive 2010/63/EU).

### Mitochondrial Isolation

Mitochondria were isolated from freshly excised mouse liver using a standard differential centrifugation protocol. The liver was rinsed in HIM buffer supplemented with 0.2% fatty acid-free BSA (Cat. No. A7030, Sigma-Aldrich). HIM buffer was prepared with the following composition per 1 liter: 200 mM mannitol (40 g, Cat. No. M4125, Sigma-Aldrich), 70 mM sucrose (23.95 g, Cat. No. S7903, Sigma-Aldrich), 10 mM HEPES (2.38 g, Cat. No. H3375, Sigma-Aldrich), 1 mM EGTA (0.38 g, Cat. No. E3889, Sigma-Aldrich), and 1 mM MgClL (Cat. No. M8266, Sigma-Aldrich), with pH adjusted to 7.5 using KOH (Cat. No. P1767, Sigma-Aldrich).

For mitochondrial isolation, the liver was placed into a Petri dish with 3 mL of HIM buffer containing 0.2% BSA, minced into small pieces, and homogenized with 6–8 strokes using a manual 7 mL glass homogenizer (Cat. No. 885300-0007, Kimble Chase). The homogenate was diluted with 7.5 mL of HIM buffer and centrifuged at 600 g for 10 minutes using a Sorvall WX 100+ centrifuge with SS-34 rotor. The supernatant was transferred to a clean tube, and the pellet was discarded. The supernatant was then centrifuged at 7000 g for 15 minutes. The resulting mitochondrial pellet was resuspended in HIM buffer, centrifuged at 700 g for 10 minutes to remove remaining cell debris, and the supernatant was subjected to a final centrifugation at 7000 g for 15 minutes. The purified mitochondrial pellet was resuspended in 200 µL of HIM buffer for downstream applications.

### Atomic Force Microscopy (AFM)

For AFM, Nanosensors qp-BioAC-CI-50 probes (Cat. No. CB2, Nanosensors) were used, specifically the CB2 probe with a resonance frequency in buffer of approximately 15 kHz, nominal stiffness of 0.1 N/m, and apex radius of 30 nm. This probe was used in tapping mode for imaging mitochondrial morphology and for elasticity measurements. Two different AFM systems were utilized: Bruker Multimode AFM was used for overall imaging and power spectra in tapping mode. To record power spectra, the AFM tip was placed on selected mitochondria without lateral scanning, and the z-piezo signal (height in volts, approximate conversion factor 0.2 nm/mV) was routed to a Picoscope 3205F digital oscilloscope to acquire data for 10 seconds at a rate of 10 kS/s. Height signal was collected in volts. Elastic modulus and height measurements were conducted using the JPK Nanowizard III system in QI mode, which collects force versus distance curves at each pixel, with the maximum applied force set to 150 pN. 5×5 µm^2^ images were captured at 128×128 pixel resolution.

Samples were prepared by immersing mica substrates in a poly-L-lysine (PLL) solution (Cat. No. P8920, Sigma-Aldrich) for 24 hours prior to the experiment, allowing PLL to adsorb onto the surface. Isolated mitochondria suspended in buffer were then deposited onto the coated substrates, incubated for 10 minutes to facilitate adhesion, and washed four times with 200 µL of buffer to remove non-adherent particles.

A few representative scans of fixed and dried mitochondria were acquired using a Bruker Multimode in Tapping mode with a 240AC-NA probe (OPUS) (such as that shown in Figures S1D, E) to confirm mitochondrial structural identity at higher image quality. Prior to imaging, the samples were fixed with 0.2% glutaraldehyde solution (Cat. No. G5882, Sigma-Aldrich) for 15 minutes, a standard procedure to preserve ultrastructure, followed by drying for atomic force microscopy (AFM).

AFM image processing was conducted using JPK SPM Data Processing software and Gwyddion^45^. Young’s modulus was analyzed at the center of each mitochondrion. Statistical analyses including ANOVA and graphical outputs were generated using GraphPad Prism (Cat. No. PRISM9, GraphPad Software). Fast Fourier Transform (FFT) analysis was performed using OriginLab (Cat. No. ORG-2024, OriginLab Corporation).

To aid visual clarity, the power spectra were averaged over intervals of 1 Hz and the spectra were normalized to the average value of the featureless region between 13-20 Hz. The units in these spectra are thus reported as arbitrary units and are meant to accentuate the finer spectral features (or lack thereof). The integrated power spectra are obtained by integrating their value over the entire DC to 20Hz range.

### Transmission Electron Microscopy (TEM)

MLM were fixed with 4% paraformaldehyde (Cat. No. 15710, Electron Microscopy Sciences) and 2% glutaraldehyde (Cat. No. 16210, Electron Microscopy Sciences) in 0.1 M cacodylate buffer (5 mM CaClL, pH 7.4) for 24 hours. Samples were then post-fixed in 1% osmium tetroxide (Cat. No. 19140, Electron Microscopy Sciences) with 0.5% potassium hexacyanoferrate (Cat. No. 1049730100, MilliporeSigma) and potassium dichromate (Cat. No. 1048650500, MilliporeSigma) for 1 hour. Staining was performed with 2% uranyl acetate (Cat. No. 22400, Electron Microscopy Sciences) in double-distilled water for 1 hour. Samples were dehydrated through a graded ethanol series and embedded in epoxy resin (Cat. No. AGR1045, Agar Scientific).

Ultrathin sections of 70 nm were prepared using a Leica EMUC7 ultramicrotome and collected on 200-mesh copper grids. The sections were stained with lead citrate (Cat. No. 15326, Sigma-Aldrich) and imaged using a Tecnai T12 Spirit transmission electron microscope (Thermo Fisher Scientific). Digital micrographs were acquired using a Gatan OneView camera.

### Respirometry assays

#### Oxygen Consumption of Isolated Liver Mitochondria

Oxygen consumption of isolated mitochondria was assessed using the Seahorse Bioscience XF96 platform (Agilent Technologies, USA), as described by Agilent43,44. Mitochondria were isolated as described above in the mitochondrial Isolation section, the isolated mitochondria were seeded at a density of 2.5ug per well in a Seahorse microplate on the day of the experiment. The mitochondria final pellet was resuspended in a minimal volume of MAS+BSA. Total protein (mg/mL) was determined using the Bradford Assay reagent (Bio-Rad). To minimize variability between wells, mitochondria were first diluted 10x in cold 1x MAS + substrate, then subsequently diluted to the needed concentration required for plating. Note that substrate is included in the initial dilution and is present during the centrifugation step. Next, 50 µL of mitochondrial suspension was delivered to each well (except for background correction wells) while the microplate plate was on ice. The Seahorse XF96 Cell Culture Microplate was then transferred to a centrifuge equipped with a swinging bucket microplate adaptor, and spun at 2,000 g for 10 minutes at 4°C. After centrifugation, 130 µL of prewarmed (37°C) 1x MAS + substrate (different combinations of substrates were used for the activation of different complexes) was added to each well. The mitochondria were viewed briefly under a microscope at 20x to ensure consistent adherence to the well. The plate was then transferred to the Agilent Seahorse XFe/XF96 Analyzer, and the experiment initiated. Different combinations of injections were used for the activation or inhibition for different complexes. The substrates and inhibitors we used in injections are as follows: ADP (4 mM final); FCCP (4 µM, final; Antimycin A (4 µM, final). Rotenone (0.5 µM, final).

#### Reagents and Solutions

The following reagents were freshly prepared and added under controlled conditions: Oxidative phosphorylation was stimulated by the addition of ADP at a final concentration of 200 µM (Cat. No. A2754, Sigma-Aldrich), which promotes ATP synthesis and activates respiration. To inhibit Complex I, rotenone was used at a concentration of 1.5 µM (Cat. No. R8875, Sigma-Aldrich), while FCCP at 1 µM (Cat. No. C2920, Sigma-Aldrich) was applied to uncouple mitochondrial respiration. Malate (5 mM, Cat. No. M1000, Sigma-Aldrich) and glutamate (10 mM, Cat. No. G5889, Sigma-Aldrich) were used as Complex I substrates, whereas succinate at 10 mM (Cat. No. S2378, Sigma-Aldrich) was used to activate Complex II.

To inhibit Complex III, antimycin A was applied at a concentration of 1 µM (Cat. No. A8674, Sigma-Aldrich). Valinomycin (1.8 µM, Cat. No. V1644, Thermo Fisher Scientific) was used as a potassium ionophore to increase membrane permeability. The electron donor TMPD was used at 0.5 mM (Cat. No. 87890, Sigma-Aldrich), and its oxidation was enhanced with ascorbate (10 mM, Cat. No. A92902, Sigma-Aldrich). Complex IV was inhibited with sodium azide at 5 mM (Cat. No. S2002, Sigma-Aldrich). ΔΨ_m_ was assessed using the fluorescent dyes JC-1 (5 µM, Cat. No. T3168, Thermo Fisher Scientific) and TMRM (0.5 µM, Cat. No. T668, Thermo Fisher Scientific). Mitochondrial visualization was performed with MitoTracker Green FM at a concentration of 0.5 µM (Cat. No. M7514, Thermo Fisher Scientific).

#### Image Processing and Statistical Analysis

All statistical analyses were performed using GraphPad Prism. Data was analyzed for significance using ANOVA tests where applicable.

#### Figure Preparation

Figures were processed and annotated using software tools including BioRender and digital microscopy platforms.

### Supporting Information Available

Additional experimental details, confocal and TEM images, AFM topography maps, oxygen consumption rate data, and frequency-dependent power spectra (PDF).

## Acknowledgments

We would like to express our deepest gratitude to the following individuals and groups for their invaluable contributions to this work. We thank Dr. Ziv Porat (Department of Chemical Research Support, Weizmann Institute of Science) for his assistance with flow cytometry (FACS). We are grateful to Dr. Elena Ainbinder and Dr. Kira Orlovsky (SeaHorse Facility, Harry Levine Family Building, Weizmann Institute of Science) for their technical support with mitochondrial respiration assays. Special thanks to members of the Gross research group for their scientific guidance throughout this study. We also extend our heartfelt appreciation to Dr. Inna Grosheva for her valuable input and support. We acknowledge Nili Dezorella and Smadar Zaidman (Department of Chemical Research Support, Weizmann Institute of Science) for their expert assistance with electron microscopy. Finally, we thank Dr. Melanie Bokstad Horev (Department of Biomolecular Sciences and Department of Immunology and Regenerative Biology, Imaging Center, Weizmann Institute of Science) for her support with imaging analyses.

