## Supplemental Figure 1 and 2 for "Non-Invasive Mechanical-Functional Analysis of Individual Liver Mitochondria by Atomic Force Microscopy"

**Supporting Information**

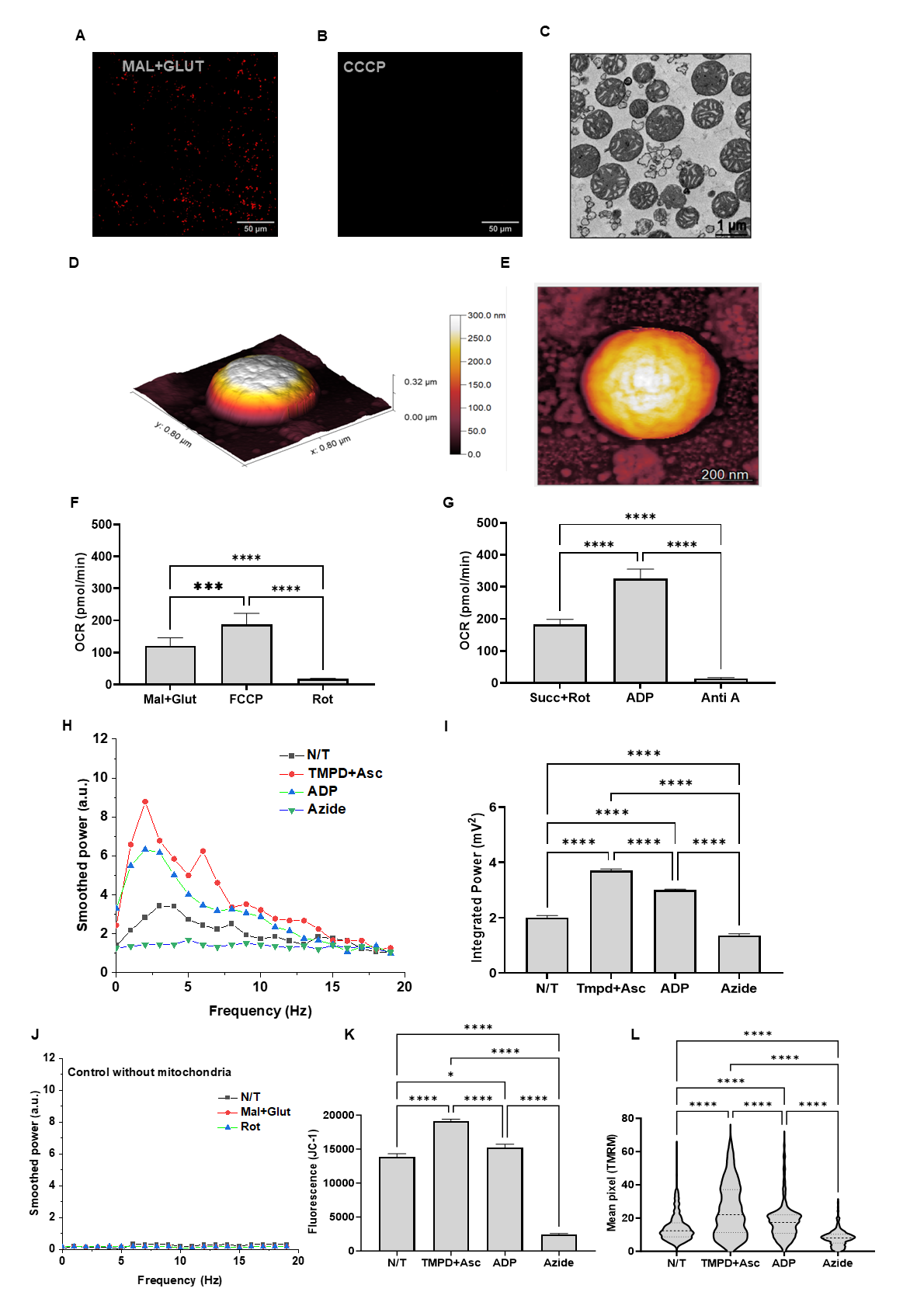

**Figure S1. Visualization and functional assessment of isolated mitochondria using confocal microscopy, TEM, AFM, and Seahorse analysis**

(A, B) Confocal fluorescence images of isolated mitochondria stained with TMRM, showing mitochondrial membrane potential (ΔΨm) under:

(A) Active conditions (malate + glutamate),

(B) Collapsed potential following CCCP treatment. Scale bars: 50 μm.

(C) Representative TEM image demonstrating intact mitochondrial ultrastructure with well-defined cristae. Scale bar: 1 μm.

(D, E) AFM topography of chemically fixed mitochondria adsorbed on poly-L-lysine–coated mica:

(D) 3D rendering of a single mitochondrion in air (tapping mode),

(E) 2D height map of the same mitochondrion. Scale bar: 200 nm.

(F, G) Oxygen consumption rates (OCR) of isolated mitochondria measured via Seahorse XF analyzer under:

(F) Complex I-linked conditions (malate + glutamate, FCCP, rotenone),

(G) Complex II-linked conditions (succinate + rotenone, ADP, antimycin A).

(H) Frequency-dependent integrated power spectra of mitochondrial height fluctuations under Complex IV-linked conditions (TMPD + ascorbate, ADP, azide), measured by AFM.

(I) Quantification of integrated power values (0–20 Hz) from spectra in panel H.

(J) Integrated power spectra in additive-only samples lacking mitochondria (negative control).

(K) JC-1 fluorescence measured by plate reader, and

(L) TMRM fluorescence measured by flow cytometry, showing ΔΨm under conditions from panel H.

All AFM analyses were performed on n = 30 mitochondria per condition. Data are presented as mean ± SEM. Statistical significance was determined using one-way ANOVA with multiple comparisons: ns = not significant; *p ≤ 0.05, **p ≤ 0.01, ***p ≤ 0.001, ****p ≤ 0.0001.

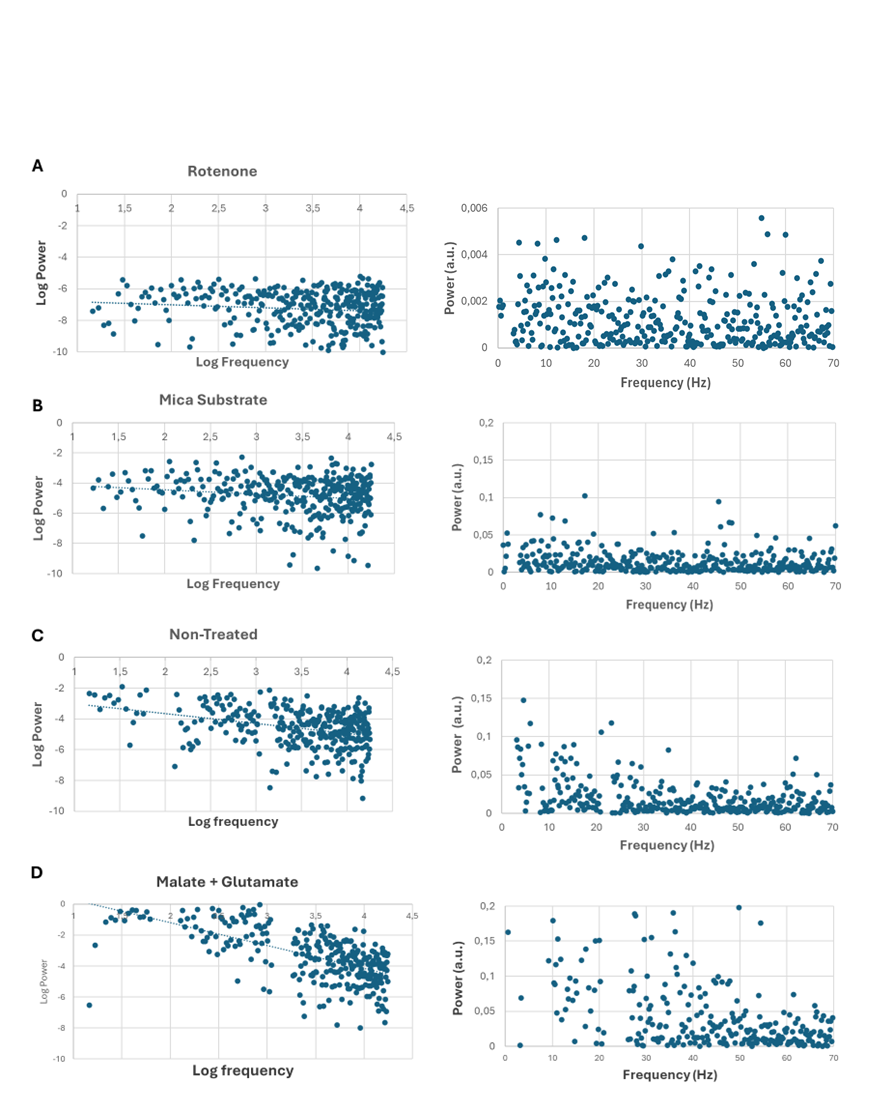

**Figure S2.** **Analysis of data for 1/fα dependence, by linear fits made to plots of log(power) vs. log(frequency), excluding sharp peaks at discrete frequency ranges**

Trend line shown on log-log plots as small dots. Plots made over range 3 – 70 Hz, α = 300

(A) Rotenone-treated α = 0.19, R2 = 0.012, p = 0.03

(B) Mica Substrate measured adjacent to mitochondria. α = 0.28, R2 = 0.026, p = 0.003

(C) Untreated mitochondria, α = 0.63, R2 = 0.119, p<0.0001

(D) Malate + Glutamate treated α = 1.5, R2 = 0.39, p<0.0001
